# p53 promotes revival stem cells in the regenerating intestine after severe radiation injury

**DOI:** 10.1101/2023.04.27.538576

**Authors:** Clara Morral, Arshad Ayyaz, Hsuan-Cheng Kuo, Mardi Fink, Ioannis Verginadis, Andrea R. Daniel, Danielle N. Burner, Lucy M. Driver, Sloane Satow, Stephanie Hasapis, Reem Ghinnagow, Lixia Luo, Yan Ma, Laura D. Attardi, Costas Koumenis, Andy J Minn, Jeffrey L. Wrana, Chang-Lung Lee, David G. Kirsch

## Abstract

Ionizing radiation induces cell death in the gastrointestinal (GI) epithelium by activating p53. However, p53 also prevents animal lethality caused by radiation-induced GI injury. Through single-cell RNA-sequencing of the irradiated mouse intestine, we find that p53 target genes are specifically enriched in stem cells of the regenerating epithelium, including revival stem cells that promote animal survival after GI damage. Accordingly, in mice with p53 deleted specifically in the GI epithelium, ionizing radiation fails to induce revival stem cells. Using intestinal organoids, we show that transient p53 expression is required for the induction of revival stem cells that is controlled by an Mdm2-mediated negative feedback loop. These results suggest that p53 suppresses severe radiation-indued GI injury by promoting intestinal epithelial cell reprogramming.

**One-Sentence Summary:** After severe radiation injury to the intestine, transient p53 activity induces revival stem cells to promote regeneration.

---

Radiation tolerance of the gastrointestinal (GI) tract limits the effectiveness of radiation therapy for thoracic, pelvic, or abdominal malignancies, such as pancreatic cancer. The radiation tolerance of the GI tract can also be exceeded in radiation accidents at nuclear power plants or after the detonation of a nuclear weapon or radioactive bomb, which can cause a lethal radiation-induced GI syndrome ^1^. There are currently no treatments to mitigate radiation-induced GI syndrome that have been approved by the US Food and Drug Administration ^1^. Thus, there is a critical need to understand the mechanisms of radiation-induced GI injury and regeneration.

During homeostasis, epithelial cell turnover in the intestine is maintained by Lgr5+ crypt base columnar (CBC) cells located at the bottom of intestinal crypts, which self-renew and give rise to differentiated progeny that constitute the intestine ^2^. The intestinal epithelium can fully recover from acute injury after a single radiation dose that ablates the Lgr5+ CBCs. However, continuous Lgr5+ CBC depletion leads to crypt loss and subsequent regeneration failure ^3, 4^, indicating that Lgr5+ CBCs must be replenished after injury to repair the intestinal epithelium. Accordingly, recent studies support a novel paradigm in which the progeny of Lgr5+ CBCs after chemical and radiation injury undergo de-differentiation to reconstitute fresh Lgr5+ CBCs and regenerate the epithelium ^5–7^.

To investigate novel signaling pathways that control the reversion of differentiated intestinal epithelial cells into stem cells following severe radiation injury, we focus on the tumor suppressor p53. We previously showed that p53 functions in the GI epithelium to prevent radiation-induced GI syndrome independent of apoptosis ^8^; however, the underlying mechanism of this phenomenon remains incompletely understood. To examine how the loss of p53 affects cell states in the irradiated intestine, we performed single-cell (sc) RNA-seq on crypt cells of the small intestines from mice that either retained functional p53 (*Villin-Cre; p53^FL/+^*) or completely deleted p53 (*Villin-Cre; p53^FL/-^)* in the intestinal epithelium 2 days after 12 Gy sub-total-body irradiation (SBI) or sham irradiation. We first generated a reference map of the transcriptional response of the regenerating intestine with intact p53. We integrated the scRNA-seq data from mice harboring wild type p53 (*Villin-Cre; p53^FL/+^*) unirradiated or at 2 days after irradiation with our previous scRNA-seq crypt data generated from wild type mice unirradiated or at 3 days after 12 Gy irradiation ^5^ (**Fig. 1A**). Unsupervised graph clustering identified 17 distinct groups of cells with different transcriptional profiles (**Fig. S1A** and **Supplementary Table 1**). Each cluster was associated with a distinct intestinal cell type based on the expression of lineage-specific gene markers (**Fig. 1B** and **Fig. S1B**). Thus, enteroendocrine, tuft cells, and Paneth cells were identified as distinct clusters by the expression of *ChgA*, *Dclk1,* and *Defa22*, respectively (**Fig. S1B**), while absorptive enterocytes clusters were identified by the expression of *Krt20* and *Alpi* and were divided into four distinct subclusters (Z1, Z3, Z4 and Z5) based on the expression of a previously described villus zonation gene signature (**Fig. S1C** and **S1D**) ^9^. We also identified two distinct clusters of crypt base columnar cells (CBC1 and 2) that expressed canonical stem cell genes *Lgr5*, *Smoc2,* and *Ascl2*, and an adjacent cluster of transit amplifying (TA) cells marked by high expression of cell cycle genes (**Fig. S1B**). Finally, we identified two additional clusters, clusters 10 and 14, both of which lacked expression of intestinal differentiation markers and canonical stem cell genes (**Fig. S1B**). Remarkably, these clusters were absent in unirradiated intestines, but they emerged and expanded on days 2 and 3 after irradiation (**Fig. 1C**).

**Fig. 1.**
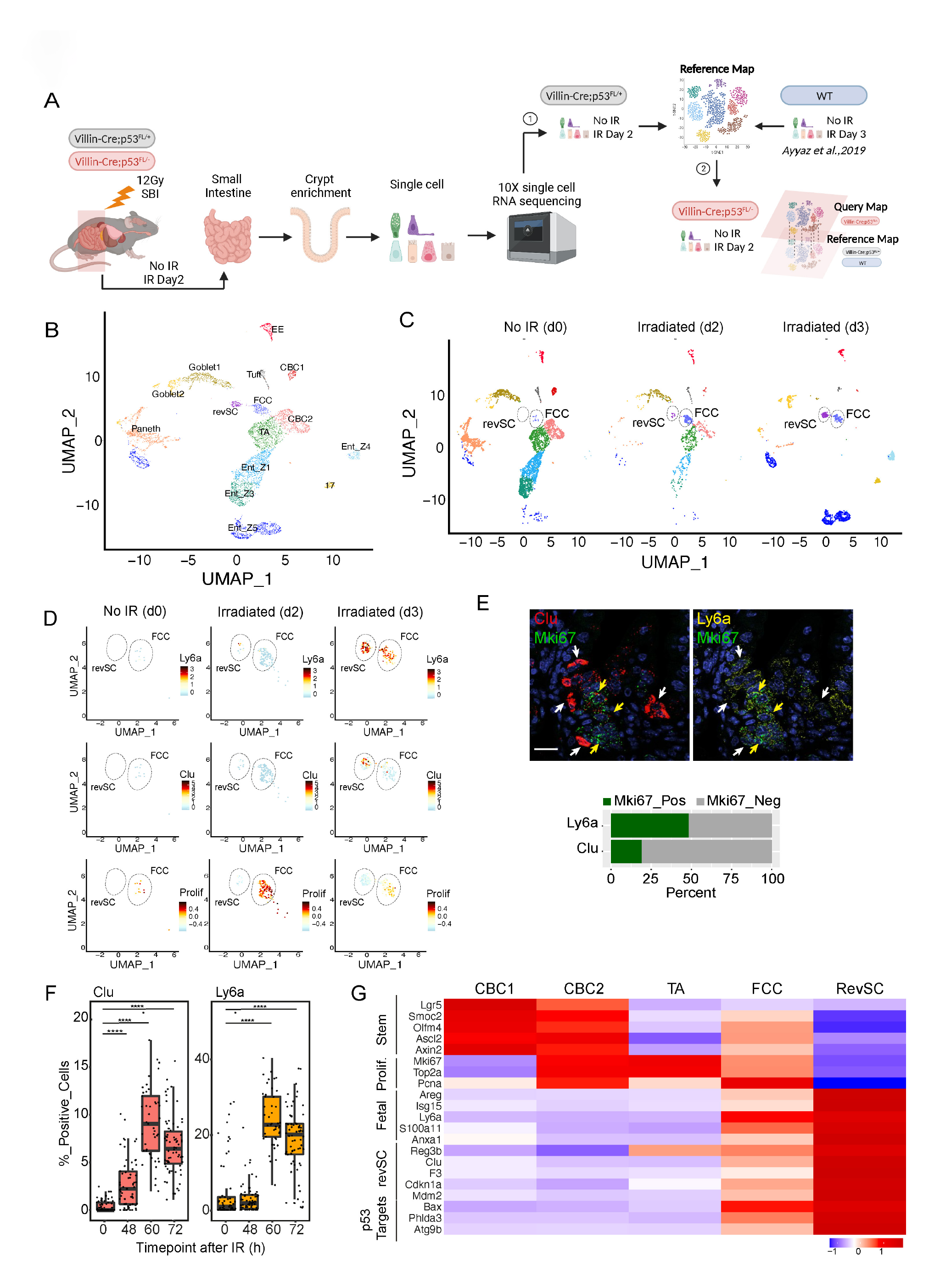
Revival Stem Cells and Fetal-like CBC cells are induced after irradiation and express p53 transcriptional targets. **(A)** Schematic indicating workflow used to profile the transcriptome of the regenerating intestinal epithelium at single-cell resolution and generation of a reference and query map. The reference map was generated using intestinal epithelial cells from p53 WT mice (p53^FL/+^ and p53 WT) non-irradiated (d0), two (d2) and three (d3) days after 12 Gy SBI (step 1). Next, the reference map was used to query the transcriptional profile of intestinal epithelial cells from p53^FL/-^ mice non-irradiated (d0) and two days (d2) after 12 Gy SBI (step 2). **(B)** UMAP representation of the reference map including all samples (non-irradiated, two and three days after 12 Gy of SBI). Intestinal epithelial cell clusters are labeled based on their transcriptional profile. **(C)** UMAP plots representing changes in cellular clusters between non-irradiated, two (d2) and three (d3) days after 12 Gy of SBI. Dashed lines highlight revival stem cells (revSC) and fetal-like CBC cells (FCC) clusters that emerge and expand only after radiation exposure. **(D)** UMAP plots showing the mRNA expression pattern of *Ly6a* and *Clu* transcripts and the proliferation gene signature in revSC and FCC clusters in non-irradiated, 2 and 3 days after irradiation. **(E)** Representative sm-FISH images of *Clu* (left) or *Ly6a* (right) and *Mki67* mRNA molecules expressed in intestinal tissue sections from mice three days after irradiation. Most of *Clu*+ cells are *Mki67*- (white arrowheads) whereas 50% of *Ly6a*+ cells are also *Mki67+* (yellow arrowheads). Quantification of double *Mki67+ Ly6a+* and *Mki67+Clu+* cells in intestinal tissue sections from mice 3 days after irradiation. Scale bar=25μm. **(F)** Percentage of *Clu* and *Ly6a* positive cells in intestinal tissue sections from non-irradiated mice (0h) or 48h, 60h and 72h after 12 Gy of SBI. Each dot represents a single tissue area quantified. (n= 3-6 mice/condition, for each mouse n= 15-20 intestinal tissue areas). ****p-val<0.0001 calculated using unpaired t-test. **(G)** Heatmap showing that p53 transcriptional targets are specifically expressed in revSCs and FCCs, but not in LGR5+ crypt base columnar cells (CBC1 and CBC2) or transit amplifying (TA) cells.

Regeneration of the intestine is associated with the induction of Yap/Taz signaling and a fetal-like transcriptional program, which can be identified by the induction of Stem Cell Antigen-1 (Sca1), also called Ly6a, in regenerative cells of the injured intestinal epithelium ^10, 11^. We found that Ly6a expression was detected in these two clusters of undifferentiated cells (clusters 10 and 14) that emerged after irradiation (**Fig. 1D, S1B** and **S1E**). Next, we investigated whether these clusters were also associated with the gene signature of revival stem cells (revSCs), a unique population of cells that is de-differentiated from recent progeny of Lgr5+ CBCs through a Yap-dependent reprogramming process and can regenerate the intestinal epithelium in response to radiation injury ^5, 12^. We found that only cells in cluster 14 expressed revSC-associated genes such as *Clu*, *Areg*, *Anxa2* (**Fig. 1D, S1B,** and **S1E**). We confirmed these observations by single molecule fluorescence in situ hybridization (smFISH) in the intestinal tissues of C57BL/6 mice exposed to 12 Gy irradiation, which showed that Ly6a expression was induced in both Clu-positive and Clu-negative cells (**Fig. 1E and S1H**). We also found that the revSC cluster lacked the expression of proliferative makers as reported previously ^5^ (**Fig. 1D** and **S1G**). smFISH assays performed in intestinal tissues at 3 days after 12 Gy further confirmed that expression of the cell cycle marker gene *Mki67* was markedly higher in Ly6a+ cells compared to Clu+ revSCs (∼50% versus 20%, respectively; **Fig. 1E**). Together, these results show that the reprogramming of the intestinal epithelium after irradiation contains two distinct populations of undifferentiated cells that are distinguished by their proliferative status: Ly6a+; Clu-proliferating cells and Ly6a+; Clu+ quiescent revSCs. Because undifferentiated Clu-proliferating cells also expressed fetal-like marker Ly6a, we named these cells fetal-like CBC cells (FCCs).

Importantly, crypts that retained p53 expression displayed a time-dependent decrease in CBCs (CBC1 and 2) with a reciprocal increase in revSCs and FCCs at 2 and 3 days after 12 Gy irradiation (**Fig. 1C** and **S2F**). Consistent with this observation, smFISH assays for Clu and Ly6a mRNA in the small intestines of C57BL/6 mice following 12 Gy showed that the percentage of either Clu+ or Ly6a+ epithelial cells were significantly increased at 60 and 72 hours after irradiation compared to unirradiated controls (0 hour) (**Fig. 1F** and **S1I**). Remarkably, p53 transcriptional targets including *Cdkn1a*, which encodes p21, *Mdm2*, *Bax*, and *Phlda3* ^13^ were expressed in revSCs and FCCs, in contrast to the low expression observed in CBCs and transit amplifying cells (TAs) (**Fig. 1G** and **Fig. S1J**). This led us to speculate that p53 may play a crucial role in regulating revSCs and FCCs during intestinal regeneration following severe radiation injury.

To assess how p53 loss affects the intestinal epithelial response to severe radiation damage, we next performed single cell profiling on GI epithelium lacking p53 from *Villin-Cre; p53^FL/-^* mice at 2 days after 0 or 12 Gy. As expected, induction of p53 transcriptional targets was abrogated in p53 mutants after irradiation (**Fig. S2A**). We then used our reference map to query the cell clusters of *Villin-Cre; p53^FL/-^* mice at 2 days after 0 or 12 Gy (**Figure 2A**). Strikingly, in contrast to mice that retained p53 (p53 intact group), the emergence of revSCs and FCCs 48 hours after 12 Gy was impaired in mice lacking p53 in the GI epithelium (p53 deleted group) (**Fig. 2B**). These results were validated using smFISH that showed Clu+ and Ly6a+ cells were significantly decreased in *Villin-Cre; p53^FL/-^* mice compared with *Villin-Cre; p53^FL/+^* mice 60 hours after 12 Gy (**Fig. 2C** and **D**).

**Fig. 2.**
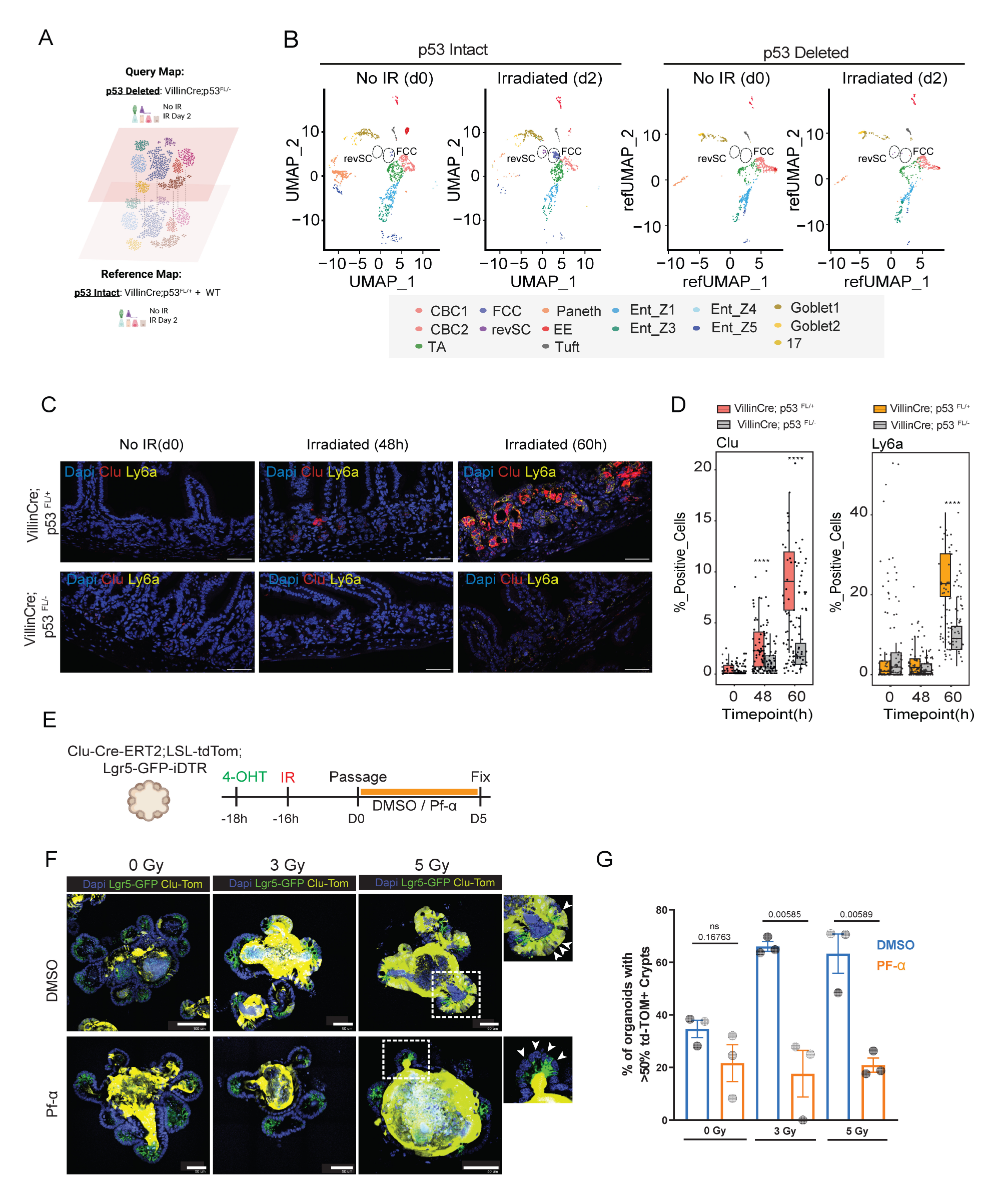
p53 is required to induce revival stem cells and fetal-like CBC cells during intestinal regeneration. **(A)** Schematic indicating the generation of query map (corresponding to p53 deleted: *Villin-Cre; p53^FL/-^*) compared with the reference map (corresponding to p53 intact: *Villin-Cre; p53^FL/+^* and WT). **(B)** UMAP plots showing changes in the cellular composition in non-irradiated (d0) or two days (d2) after 12 Gy SBI in mice with p53 WT (*Villin-Cre:p53 ^FL/+^*) or p53 KO (*Villin-Cre;p53 ^FL/-^*) intestinal epithelium. Dashed lines highlight the revSC and FCC cell clusters. **(C)** Representative images from single-molecule fluorescence in situ hybridization (sm-FISH) of *Clu* and *Ly6a* expression in mice with p53 WT (*Villin-Cre; p53^FL/+^*) or p53 KO (*Villin-Cre; p53^FL/-^*) intestinal epithelium in non-irradiated (d0) or 48h and 60h after irradiation. Scale bar=50μm. **(D)** Percentage of *Clu* and *Ly6a* positive cells quantified from (C). Each dot represents single tissue area quantified (n=3-6 mice/condition and n=15-20 intestinal tissue areas/mouse). ****p-val<0.0001 calculated using unpaired t-test. **(E)** Schematic indicating that *Clu-Cre-ERT2; LSL-tdTomato; Lgr5-GFP-iDTR* small intestinal organoids were treated with 4-hydroxytamoxifen (4-OHT) prior to irradiation (3 or 5 Gy) to induce permanent labelling of Clu+ cells and linage trace this population during organoid regeneration in control (DMSO) conditions or during p53-inhibition by Pifithrin-α (Pf-α) treatment. **(F)** Representative immunofluorescence images of organoids indicating Clu and Clu progeny (Yellow) and Lgr5 (green) on the 5^th^ day of regeneration, grown in ENR media containing either DMSO or Pf-α. White arrows indicate the absence of Clu+ cells and progeny in the crypts of irradiated organoids treated with Pf-α. Scale bar=100 and 50μm. **(G)** Quantification of the percent of organoids displaying lineage-traced crypts following the 5 day regeneration period. Each dot represents an individual experiment (N=3 experiments), with the number of organoids counted per experiment and per conditions ranging from 17-106 (n=17- 106). Error bars indicate SEM and significance was calculated via 2-tailed t-test.

Our results suggest that p53 is required for key cell state transitions underlying intestinal regeneration, in particular induction of Clu+ revSCs in the regenerating intestinal epithelium after irradiation. Therefore, we next tested whether Clu+ revSCs are implicated in intestinal regeneration at higher radiation doses that cause GI syndrome by performing lineage tracing in previously described *Clu^Cre-ERT^*^2^*^/+^*; *Rosa26^LSL-tdTomato/+^* mice (*4*). After 10 Gy or 14 Gy SBI, Clu+ revSCs and their progeny were labelled with tdTomato by administering two doses of tamoxifen (TAM) at 1 and 2 days after irradiation (**Fig. S2B**). In this mouse strain, 10 Gy and 14 Gy SBI are approximately LD_20/10_ and LD_50/10_ doses for the radiation-induced GI syndrome, respectively (**Fig. S2C)**. Of note, we observed a significant increase in the percentage of Clu-labeled regenerated intestinal crypts after 10 Gy and 14 Gy compared to 0 Gy. In addition, crypts reconstituted by Clu+ cells displayed a dose dependent increase as radiation damage increased (50% after 14 Gy vs 25% after 10 Gy) (**Fig. S2D** and **S2E**). These results demonstrate that p53- dependent induction of regenerative cell states promotes reconstitution of the small intestinal crypts after high-dose irradiation that causes the GI syndrome, which is consistent with our previous work ^5^.

To investigate the impact of acute p53 inhibition on the regeneration of intestinal crypts via Clu+ revSCs, we established an ionizing radiation (IR) damage model in intestinal organoids from *Clu^Cre-ERT^*^2^*^/+^*; *Rosa26^LSL-tdTomato/+^*; *Lgr5^DTR-GFP^* mice to enable *in vitro* lineage tracing experiments. 4-hydroxy tamoxifen (4-OHT)-treated organoids were irradiated with 0, 3 or 5 Gy followed by treatment with either vehicle alone (DMSO) or pifithrin-alpha (PF-a), a small molecule inhibitor of p53 ^14^ (**Fig. 2E**). In untreated organoids, Lgr5+ crypts derived from Clu+ offspring were 38%, but rose to >60% after 3 or 5 Gy IR (**Fig. 2G**). In contrast, PF-a treated organoids showed significantly decreased percentages of Lgr5+; tdTomato+ crypts after 3 and 5 Gy (**Fig. 2F** and **G**). Collectively, our findings demonstrate that loss of functional p53 impairs the emergence of revSC and FCC in the small intestine following 12 Gy *in vivo* and suppresses Clu+ revSC-dependent regeneration of crypts.

To examine the dynamics of p53 protein expression during epithelial recovery from radiation exposure, we co-stained p53 protein and Clu+ cells at 0, 1, 2, or 3 days after 12 Gy using a GFP reporter driven by the Clu promoter (Clu^GFP^) that faithfully recapitulates endogenous Clu expression in the small intestines ^5^. We observed a significant induction of nuclear p53 protein at 2 days after 12 Gy, the timepoint when Clu+ revSCs first start emerging in the irradiated intestine ^5^. However, the expression of p53 protein was transient as it was substantially diminished by 3 days after irradiation ^15^ (**Fig. 3A**, **3B** and **Fig. S3A, S3B**). The decrease in nuclear accumulation of p53 protein was associated with the induction of Mdm2, an E3 ubiquitin-protein ligase that degrades p53 ^16^ (**Fig. 3C**). Indeed, at 3 days post irradiation, *Mdm2* mRNA was substantially induced in both revSCs and FCCs, but not in CBCs and TAs (**Fig. 3D**). To investigate how the p53-Mdm2 negative feedback loop controls revSCs, we utilized the *Clu^Cre-ERT2/+^*; Rosa26^LSL-tdTomato/+^; *Lgr5^DTR-GFP^* organoid IR damage model, and performed lineage tracing after irradiation as described above, but in this case treated with Nutlin-3, a small molecule Mdm2 inhibitor ^17^ (**Fig. 3E**). Continuous treatment of the organoids with Nutlin-3 after irradiation markedly increased p53 protein levels and decreased Clu-mediated crypt regeneration following 3 and 5 Gy (**Fig. 3F**). In addition, Nutlin-3 treated irradiated organoids were smaller in size, had fewer buds, and did not survive past secondary passaging, indicating that the stem cell compartment was lost in the presence of continuous p53 activity (**Fig. 3G**). Together, our results suggest a critical role for transient p53 controlled by the p53- Mdm2 negative feedback loop in driving the Clu+ revSC regenerative pathway after severe radiation injury.

**Fig. 3.**
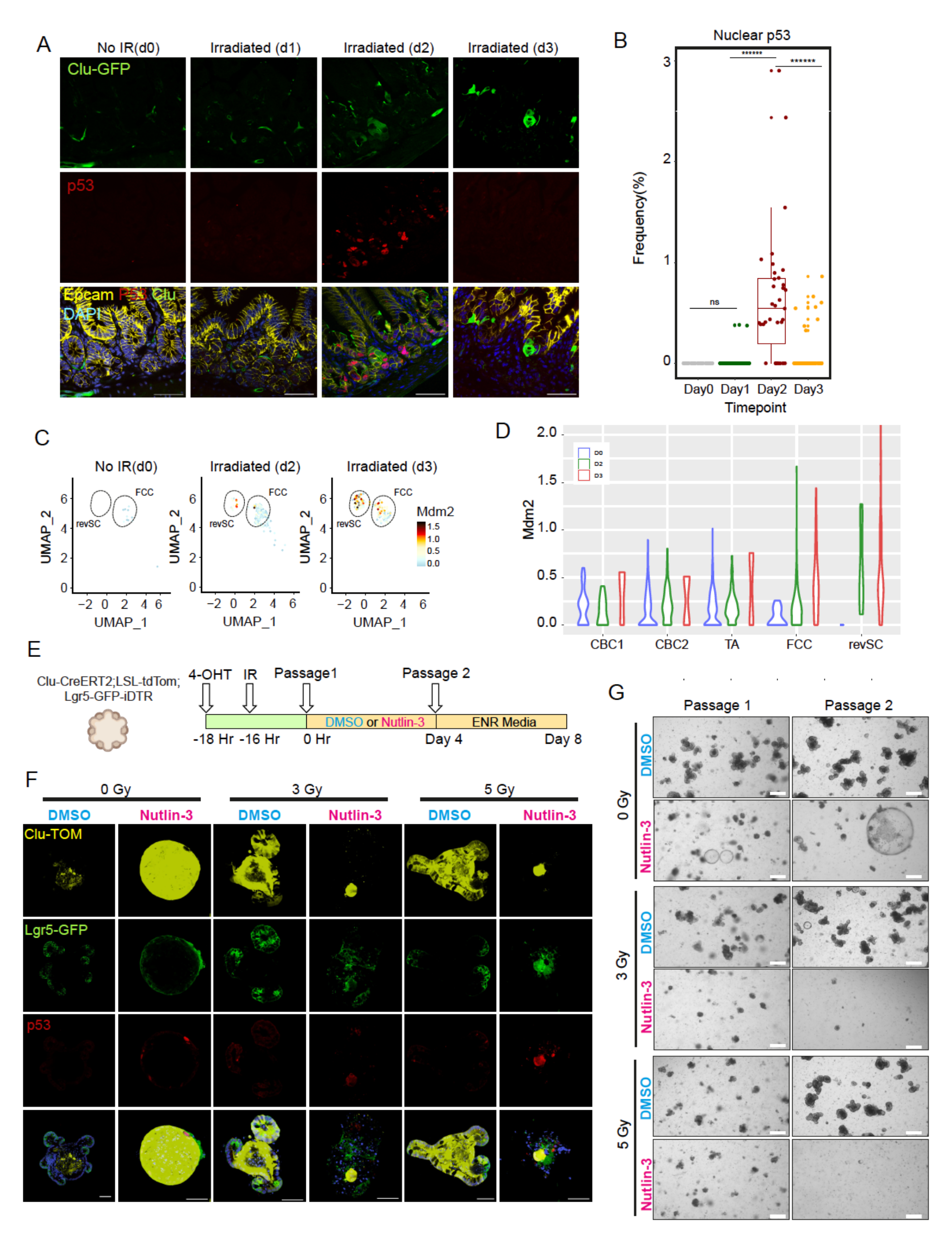
Constitutive expression of p53 impairs intestinal tissue regeneration after irradiation. **(A)** Immunofluorescence of Clu-GFP, p53 (red), and Epcam (yellow) on intestinal tissue sections 1, 2, and 3 days after 12 Gy SBI. Scale bar=50μm. **(B)** Quantification of the percentage of cells with nuclear p53 from (A). Data from only one experiment (3 mice/time point) is shown with each dot representing single cross section.****p-val<0.000001 calculated using Welch’s t-test. **(C)** UMAP plots showing the *Mdm2* mRNA expression in revSC and FCC populations at day 0, or 2 and 3 days after 12 Gy SBI. **(D)** *Mdm2 mRNA expression* in columnar crypt base (CCB1 and 2), transit amplifying (TA), fetal-like CBC cells (FCC), and revival stem cells (revSC) clusters at Day 0 or 2 and 3 after 12 Gy of SBI. **(E)** Schematic indicating that Lgr5-GFP; Clu-CreERT2 small intestinal organoids were treated with 4-OHT to induce a lineage trace form Clu+ cells before irradiation (3 or 5 Gy), all prior to passaging. Organoid regeneration was tracked over a 4-day period, with growth under control (DMSO) or p53-activated conditions (Nutlin-3) (**4F**). A second passage was then made onto normal ENR growth media (**4G**). **(F)** Representative immunofluorescence images of Clu and Clu progeny (yellow), Lgr5 (green) and p53 (red), indicating that under continuous p53 activation (Nutlin-3), organoids show an inability to recover from IR conditions by day 4 of regeneration. Scale bar=50μm. **(G)** Bright field images 1 and 2 passages after irradiation, indicating that organoids show an inability to recover from continuous p53 activation, with culture collapse evident after the initial treatments. Notably, recovery is seen in organoids placed onto media containing DMSO after irradiation. Scale bar=500μm.

To define possible mechanisms by which p53 promotes proper regeneration of irradiated intestinal epithelium, we performed gene set enrichment analysis (GSEA) to identify signaling pathways significantly enriched after irradiation in crypts from mice with and without functional p53 (**Fig. 4A**). Notably, we observed several pathways that were significantly upregulated in irradiated mice with intact p53, but they were downregulated after irradiation in mice that lost p53 in the GI epithelium. These pathways mainly involve in the control of cell proliferation (Myc targets, oxidative phosphorylation, and E2F targets) or the DNA damage response (G2/M checkpoint and DNA repair). In addition, we also observed significant upregulation of interferon alpha response after irradiation only in p53 deficient mice (**Figure 4A**). These findings suggest that p53 may function as a transcription factor to have a pleotropic impact on the regeneration of irradiated intestinal epithelium.

**Fig. 4.**
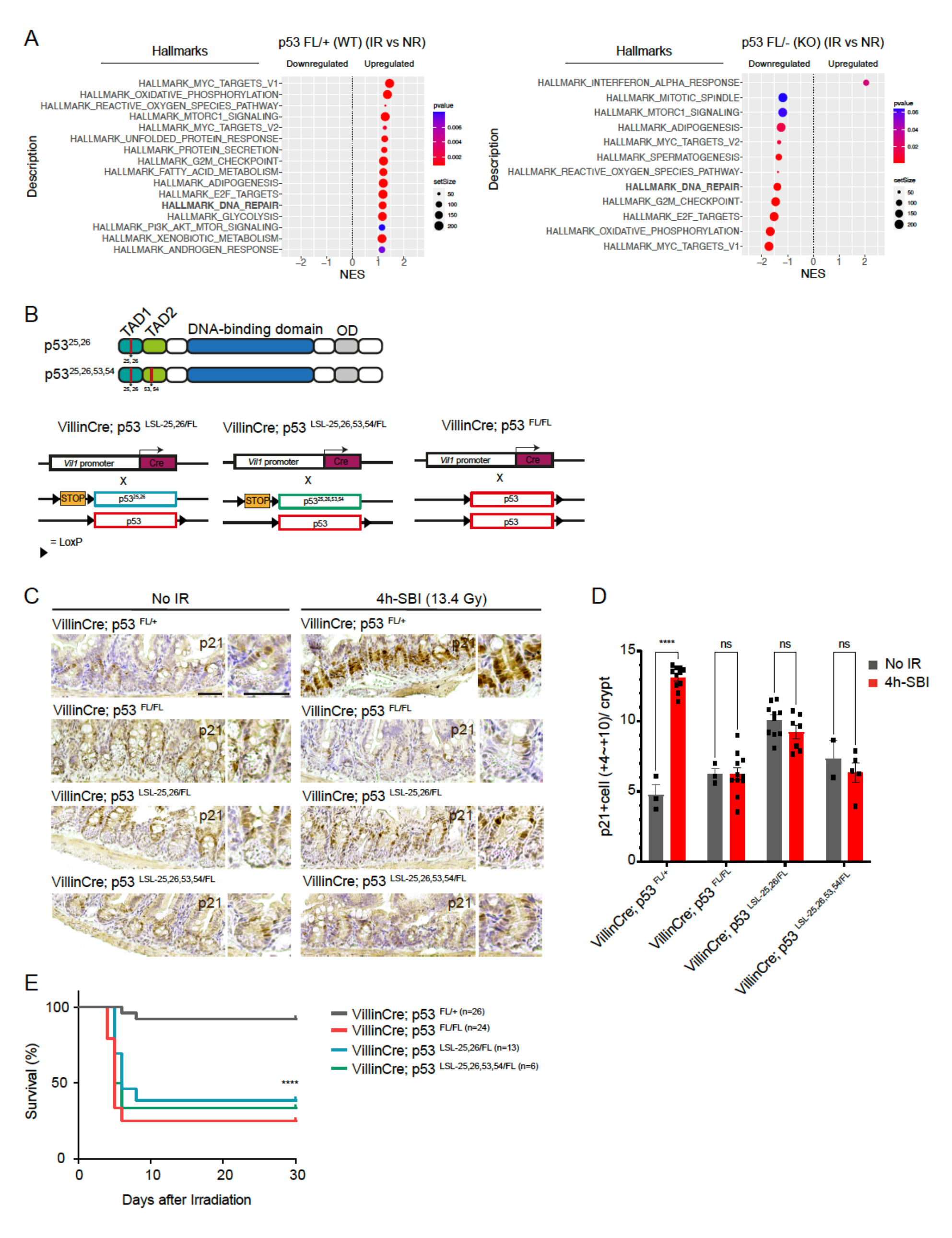
p53 TAD domains are required to prevent radiation-induced GI syndrome. **(A)** Gene set enrichment analysis (GSEA) showing top upregulated or downregulated gene sets two days after irradiation (IR) in p53 WT (FL/+) or p53 deficient (FL/-) intestinal epithelial cells. **(B)** Schematic of p53 protein domains showing the locations of the transactivation domains (TAD) mutation residues (upper) and schematic of p53 alleles of the *Villin-Cre* mice used in this study (lower). TAD1 and TAD2: transactivation domains 1 and 2. OD: oligomerization domain. **(C)** Representative images of immunohistochemistry staining of p21 protein on intestinal tissue sections from p53 WT or p53 mutants non-irradiated (No IR) or 4 hours after 13.4 Gy SBI. Scale bars=50μm. **(D)** Quantification of p21 staining from (C). p21-positive cells within the +4 to +10 region of the crypt were counted. Each dot represents one mouse. The mean is graphed with SEM. **** represents p<0.0001 in a 2-way ANOVA comparison using Bonferroni’s multiple comparison test. **(E)** Kaplan-Meier curves of WT (p53^FL/+^) or p53 mutants (p53^FL/FL^, p53^LSL-25, 26/FL^, and p53^LSL-25, 26, 53, 54/FL^) mice following 13.4 Gy of SBI (total n=69). P value is calculated using a long-rank test.

The transcriptional activity of p53 is controlled by two N-terminal transactivation domains (TAD): TAD1 and TAD2 ^13^. To dissect the role of p53-mediated transactivation in the radiation-induced GI syndrome, we conducted experiments using mice that harbor mutations in *p53* that impair either TAD1 alone or both TAD1 and TAD2. Previous studies using these mice demonstrate that the transcription of genes that affect the canonical response of p53 to acute DNA damage is mainly dependent on TAD1, whereas the transcription of genes that mediate the non-canonical response of p53 is dependent on TAD1 and/or TAD2 *in vivo* ^13^. Thus, we generated compound transgenic mice using *Villin-Cre* to delete one floxed p53 allele (FL) and remove a transcription/translation STOP cassette to activate expression of p53^25, 26^ with mutations in TAD1 alone (*Villin-Cre*; *p53^LSL-25,26/FL^*), which impairs the response of p53 to acute DNA damage. The same strategy was used to activate the expression of p53^25, 26, 53, 54^ with mutations in both TAD1 and TAD2 (*Villin-Cre*; *p53^LSL-25,26,53,54/FL^*), which impairs all p53- dependent transcriptional responses (**Fig. 4B**). The expression of the TAD domain mutants in the GI epithelial cells was confirmed by immunohistochemistry (IHC) (**Fig. S4A-D**).

To examine whether p53-dependent transcriptional programs were abrogated in mice with GI-specific p53 TAD mutant protein expression, we performed IHC on the intestine of unirradiated mice and mice 4 hours after 13.4 Gy SBI to detect the canonical p53 transcriptional target p21 ^18^. In mice that retained wild type p53 (*p53^FL/+^*), we observed induction of p21 after SBI. However, the *Villin-Cre*; *p53^LSL-25,26/FL^* and *Villin-Cre*; *p53^LSL-25,26,53,54/FL^*mice failed to induce p21 after irradiation similar to mice that lacked p53 (*Villin-Cre; p53^FL/^*^FL^) (**Fig. 4, C** and **D**). To evaluate a functional endpoint of the p53 transcriptional program, we also performed IHC for cleaved-caspase-3 to quantify radiation-induced apoptosis in the GI epithelium, which is p53- dependent ^19^. In mice that retained wild type p53 (*p53^FL/+^*), we observed a significant increase in the number of cleaved-caspase-3 positive intestinal cells after SBI, but in mice expressing p53 TAD mutants (*Villin-Cre*; *p53^LSL-25,26/FL^* or *Villin-Cre*; *p53^LSL-25,26,53,54/FL^*), the number of cleaved-caspase-3 positive intestinal cells was not significantly induced by irradiation and was similar to mice that lacked p53 (*Villin-Cre*; *p53^FL/FL^*) (**Fig. S4, E** and **F**).

Mice containing mutated GI epithelia were then treated with 13.4 Gy SBI and followed for radiation-induced GI syndrome. In contrast to mice that retained one wild type allele of p53 in the GI epithelium (*Villin-Cre; p53^FL/+^*), mice expressing p53 with mutations in TAD1 alone (*Villin-Cre*; *p53^LSL-25,26/FL^*) or in both TAD1 and TAD2 (*Villin-Cre*; *p53^LSL-25,26,53,54/FL^*) were sensitized to radiation-induced GI syndrome similar to mice that lacked p53 (*Villin-Cre; p53^FL/^*^FL^) (**Fig. 4E** and **Fig. S4G)**. These results demonstrate that a p53-dependent response to acute DNA damage is necessary to protect mice from radiation-induced GI syndrome. Taken together, our findings from scRNA-seq, intestinal organoids, and genetically engineered mice harboring different p53 TAD mutations in the GI epithelium indicate that p53-dependent production of revSCs requires the p53 transcriptional response to DNA damage to regenerate the epithelium after severe radiation damage.

In summary, our results provide novel insight into a long-standing question in radiation biology: how does the tumor suppressor p53 suppress the development of the radiation-induced GI syndrome? ^8, 20–22^ Our results indicate that transient activation of p53 after irradiation is necessary to promote the emergence of revSCs. We demonstrate that Clu+ revSCs contribute to a significant fraction of regenerating epithelial crypts during the radiation-induced GI syndrome and the revSCs have also been previously shown to increase animal survival following gut damage ^5^. Importantly, while acute inhibition of p53 impairs crypt regeneration via revSCs, prolonged p53 activation through inhibition of Mdm2 also suppresses the ability of revSCs to regenerate crypts following irradiation. These results define a key role of the p53-Mdm2 feedback loop in controlling the emergence of revSCs during intestinal regeneration following severe radiation injury.

## Supplementary Figure Legends

**Supplementary Fig. 1. Identification and transcriptional characterization of intestinal epithelial cell clusters identified by single-cell RNA sequencing.**

**(A)** UMAP

**(B)** Dotplot showing the expression of lineage-specific gene markers (columns) across all epithelial cell clusters (rows).

**(C)** Percentage of cells in each identified intestinal epithelial cell cluster from non-irradiated intestines (d0) and two (d2) and three (d3) days after 12 Gy SBI.

**(D)** Identification of differentiated cell clusters based on the expression of the pan-differentiation gene *Krt20*.

**(E)** Heatmap showing the expression of enterocyte zonation gene signatures across the Krt20+ cell clusters.

**(F)** Identification of FCC and revSC clusters based on the expression of *Ly6a* and *Clu* genes on the UMAP plot.

**(G)** Violin plot showing the expression of *Lgr5*, *Pcna*, *Ly6a* and *Clu* genes across the columnar crypt base (CCB1 and 2), transit amplifying (TA), fetal-like CBC cells (FCC) and revival stem cells (revSC) clusters.

**(H)** Representative images of smFISH assay of Ly6a and Clu gene expression in intestinal tissues 72h after 12 Gy of SBI. Inset highlights Ly6a+ Clu-and Ly6a+ Clu+ cells. Scale bars=50μm.

**(I)** Representative images of single-molecule fluorescence in situ hybridization (sm-FISH) assay of *Clu* and *Ly6a* gene expression in intestinal tissue sections from non-irradiated mice or 48h, 60h and 72h after 12 Gy SBI. Scale bars=50μm.

**(J)** UMAPS showing the expression of p53 transcriptional gene targets across the different intestinal epithelial cell types in non-irradiated (NR) and 2 and 3 days after 12 Gy SBI.

**Supplementary Fig. 2. Clu+ cells contribute to the regeneration of the small intestines following various doses of sub-total body irradiation.**

**(A)** Heatmap showing the expression of p53 transcriptional target genes in p53 WT (*Villin-Cre; p53^FL/+^*) or p53 deficient (*Villin-Cre; p53 ^FL/-^*) in non-irradiated (NR) or two days after 12 Gy SBI.

**(B)** Clu-CreER; LSL-tdTomato mice exposed to 0, 10 and 14 Gy SBI were injected with 100 mg/kg tamoxifen at 24 and 48 hours after irradiation to induce CreER-mediated gene recombination and expression of tdTomato. The small intestines were harvested from mice that survived 10 days after irradiation.

**(C)** Kaplan-Meier curves of mice following 0, 10, or 14 Gy SBI (total n=16). P value is calculated using a long-rank test.

**(D)** Representative tissue sections of the small intestine harvested 10 days after 0, 10 and 14 Gy SBI. The DAPI signal is shown in blue and the tdTomato signal is shown in red. Scale bars=500µm.

**(E)** Quantification of lineage tracing events of the small intestines harvested 10 days after irradiation. Each dot represents one mouse.

**Supplementary Fig. 3. Transient activation of p53 in intestinal epithelial cells after irradiation.**

**(A)** Representative IHC staining of p53 in *Villin-Cre; p53^FL/+^* mice at various time points post 12 Gy SBI. An example of a positively staining crypt for each timepoint is highlighted in the adjacent panel photo. Scale bars =100μm.

**(B)** Quantification of median number of p53-positive cells per crypt in *Villin-Cre; p53^FL/+^* mice at various timepoints post 12 Gy SBI.

**Supplementary Fig. 4. Kaplan-Meier curves and quantification of cleaved-caspase 3 staining in the intestinal crypts of mice after irradiation.**

**(A-D)** p53 IHC in the intestinal epithelium of (A) p53 WT (*Villin-Cre; p53^FL/+^*), (B) *Villin-Cre; p53^FL/FL^*, (C) *Villin-Cre; p53^LSL-25, 26/FL^* and (D) *Villin-Cre; p53^LSL-25, 26, 53, 54/FL^* mice. Strong p53 expression is seen in (C) and (D) because in the TAD mutants p53 does not bind to Mdm2. Therefore, p53 is overexpressed.

**(F)** Representative images of IHC of cleaved caspase-3 in intestinal tissue sections from p53 WT or p53 mutants non-irradiated (No IR) or 4 hours after 13.4 Gy SBI. Scale bar=100um.

**(G)** Quantification of caspase-3 positive cells from (C). **** represents p<0.0001 in a 2-way ANOVA comparison using Bonferroni’s multiple comparison test.

**(H)** Kaplan-Meier curves of *Villin-Cre; p53^FL/+^* or *Villin-Cre; p53^FL/FL^* in males (n=18) and female (n=14) mice following 13.4 Gy of SBI. P value is calculated using a long-rank test.

## Acknowledgments

We thank members of our laboratories for helpful feedback, Anton Berns from the Netherlands Cancer Institute for providing the p53^FL/FL^ mice, and the Cell and Animal Radiation Core Facility (RRID: SCR_022377) at the University of Pennsylvania Perelman School of Medicine.

## Funding

National Institutes of Health 2U19AI067798 (C-LL and DGK)

National Institutes of Health U19AI067773 (C-LL)

National Institutes of Health 2R35CA197616 (DGK)

National Institutes of Health 1P01CA257904 (CK, AJM, C-LL, and DGK)

Duke University School of Medicine Whitehead Scholar Award (C-LL)

The Mark Foundation for Cancer Research (AJM and CMM)

## Author Contributions

Conceptualization: H-CK, AA, CMM, AJM, C-LL, JW, DGK

Methodology: H-CK, AA, CMM, MF, LDA

Investigation: H-CK, AA, CMM, MF, IV, ARD, DNB, LD, SS, SH, RG, YM

Funding acquisition: CK, AJM, C-LL, JW, DGK

Supervision: CK, AJM, C-LL, JW, DGK

Writing – original draft: C-LL, DGK

Writing – review & editing: All authors

## Data and materials availability

All data is available in the main text or in the supplementary materials. The single-cell RNA sequencing data is available in SRA data base (PRJNA910548) upon request.

## Supplementary Materials

**Table S1.** Gene signatures used in this study to identify epithelial cell clusters.

**Table S2.** Differentially expressed genes across intestinal epithelial clusters.

## Materials and Methods

### Animal Models

All procedures with mice were approved by the Institutional Animal Care and Use Committee (IACUC) of Duke University, by the IACUC of the University of Pennsylvania and by the Canadian Council on Animal Care. All mouse strains used in this study have been described before, including *Villin-*Cre*, p53^FL/FL^*, *p53^LSL-25,26^*, *p53^LSL-25,26,53,54^*, *Clu-EGFP*, *Clu-CreERT2*, *Lgr5-CreERT2*, *Lgr5-GFP-iDTR*, Ai9, and Ai14 mice ^5,13, 23–29^. The *Villin-Cre* mice were originally obtained from the Jackson Laboratory and then bred at Duke University. The *p53^FL/FL^* mice were originally provided by A. Berns (Netherlands Cancer Institute, Amsterdam, the Netherlands) and the *p53^LSL-25,26^* and *p53^LSL-25,26,53,54^* mice were originally provided by L. Attardi (Stanford University, Stanford, CA) and then bred at Duke University. The *Lgr5-CreERT2* were obtained from Jatin Roper (Duke University). The *Clu-EGFP* mice were originally obtained from Rockefeller University (GENSAT project) and then bred at the Lunenfeld-Tanenbaum Research Institute (Toronto, ON, Canada). The *Clu-CreERT2* mice were created and bred at the Lunenfeld-Tanenbaum Research Institute, and the methods of creating the mice have been described previously ^5^. The *Lgr5-GFP-iDTR* were provided by Frederic de Sauvage (Genentech, San Francisco, CA) and bred at the Lunenfeld-Tanenbaum Research Institute. The Ai9 and Ai14 mice were obtained from Jackson Laboratory and crossed with CreERT2 mice to generate reporter mice. Mice were at least 8 weeks old at the time of irradiation. Both sexes of mice were used. Comparisons were made using littermate controls to minimize genetic differences as these experiments were performed on mixed genetic backgrounds.

### Intestinal Organoids Culture

Small intestinal organoids were cultured according to a previously described protocol established by Sato and Clevers ^30^. Crypts were collected by incubating the small intestine in PBS containing 2 mM EDTA, followed by isolation using a 70 mm cell strainer. Crypts were seeded in growth factor reduced Matrigel (BD Biosciences) and grown in Advanced DMEM/F12 (Life Technologies #12634028) supplemented with 2 mM GlutaMax (Life Technologies #35050061), 100 U/mL Penicillin/100 mg/mL Streptomycin (Life Technologies #15140122), N2 Supplement (Life Technologies #17502001), B-27 Supplement (Life Technologies #17504001), mouse recombinant Egf (Life Technologies #PMG8043), 100 ng/mL mouse recombinant Noggin (Peprotech #250-38) and 300 ng/mL human R-spondin1 (R&D Systems #4645-RS).

### Radiation Experiments

#### In vivo

For radiation-induced GI syndrome and lineage tracing experiments, 8 to 10-week old male and female mice were exposed to a single dose of SBI at Duke University. Jigs were used to hold unanesthetized mice and limit their movement. The head and the front limbs of mice were shielded by lead to prevent radiation-induced hematopoietic syndrome. SBI was performed with an X-RAD 320 biological irradiator (Precision X-ray Inc., North Branford, CT). All mice were treated 50 cm from the radiation source, and a total physical dose of 10, 13.4 or 14 Gy using X-rays of 320 kV and 12.5 mA was delivered.

For the scRNA-seq experiment, 10- to 12-week-old male and female mice were exposed to a single dose of SBI at the University of Pennsylvania. Mice were immobilized with 2.5% isoflurane anesthesia with medical air as the carrier gas (VetEquip). The head of the mouse was placed in a facemask that allows the gas to be scavenged (Xerotec Inc.). To deliver whole abdominal irradiation a variable collimator set at 20×40 mm (length x height) was used with the isocenter set at the center of the abdomen. A total physical dose of 12 Gy of X-rays using 220kV and 13mA was delivered to the mouse abdomen using the 3rd generation Small Animal Radiation Research Platform (SARRP by XStrahl).

#### In vitro

In vitro irradiation of organoids for lineage tracing experiments*, Clu-CreERT2; Rosa26-LSL-tdTomato; Lgr5-GFP-iDTR* organoids were exposed to a single dose of 3 or 5 Gy using a GammaCell 40 irradiator to induce injury 16 hours before passage.

### Lineage Tracing Experiments

To induce Cre-mediated gene recombination *in vivo*, *Clu-Cre-ERT2^+/-^; Rosa26-LSL-tdTomato^+/-^* or *Lgr5-Cre-ERT2^+/-^; Rosa26-LSL-tdTomato^+/-^* mice were administered 100mg/kg tamoxifen (Sigma #T5648) via intraperitoneal injection at time points of 24 and 48 hours after sham IR or SBI. Mice were euthanized 10 days post-SBI or sham-IR for tissue harvest. Lineage-traced crypts were quantified manually by counting crypts with ≥ 70% tdTomato signal. The percentage of lineage tracing events per slide was determined by dividing the total number of positive tdTomato crypts by the total number of crypts present. The images were quantified by two observers (S.S. and L.D.), who were blinded to the treatment. A total of 10 representative 10X images from proximal to distal were taken for each slide. The number of *Clu-Cre-ERT2^+/-^; Rosa26-LSL-tdTomato^+/-^* mice quantified were n=3 0 Gy, n=4 10 Gy, and n=4 14 Gy. The number of *Lgr5-Cre-ERT2^+/-^; Rosa26-LSL-tdTomato^+/-^* mice quantified were n=3 0 Gy, and n=1 14 Gy. The number of mice analyzed was determined by the number of mice that did not develop the radiation-induced GI syndrome prior to 10 days post-IR.

For *in vitro* lineage tracing experiments, *Clu-CreERT2; Rosa-26-LSL-tdTomato; Lgr5-GFP-iDTR* organoids were treated with 4-hydroxytamoxifen (4-OHT; Sigma #H6278; 5ug/ml) to induce a lineage trace and two hours later were irradiated as described above. At the time of passaging, organoids were grown under mock conditions, with ENR (Egf/Noggin/R-spondin) media containing DMSO, or with either Nutlin3a (Sigma #SML0580; 10 µM) to activate the p53 pathway or Cyclic Pifithrin-α hydrobromide (PF-α; Sigma #P4236; 20 µM) to inhibit the p53 pathway. Following the desired regeneration period, lineage traced crypts were quantified manually by counting the number of Lgr5-GFP+ regions that displayed ≥ 70% td-Tomato signal.

### Tissue Histology and Staining Immunohistochemistry (IHC)

Unirradiated and irradiated mice were sacrificed at indicated time points and the middle 10 to 12 cm length (jejunum) of the small intestine was isolated and flushed with cold DPBS (Gibco). The tissue was then divided into 2 cm segments for 10% formalin fixation for 16-24 hours. Prior to embedding, segments were bundled using 3M Micropore Surgical Tape (3M) for cross sectioning.

For immunohistochemistry (IHC), deparaffinized slides underwent 3% hydrogen peroxide treatment, antigen retrieval with either citrate-or Tris-based solution (Vector Laboratories), and blocking with 5% goat serum. Slides were incubated with primary antibody against p21 (ab188224; Abcam), cleaved caspase-3 (Asp175) (5A1E) (#9664; Cell Signaling Technology), or p53 (CM5; Leica). For chromogenic detection, the secondary antibodies used include Goat anti-Rabbit IgG (H+L) Cross-Adsorbed Secondary Antibody, Biotin (31822; Invitrogen). The VECTASTAIN Elite ABC system (Vector Laboratories) was applied, and 3,3’-diaminobenzidine (DAB) was used as chromogen. Slides were imaged using a Leica DM2000 LED phase contrast microscope. Cells positive for cleaved caspase-3 signals were quantified by individual crypt, and the examiner was blinded to the genotype and the treatment of the samples. Briefly, the examiner was provided with a 20X deidentified histology slide of intestinal crypts. The examiner would then determine if any intact crypts were present. Intact crypts were defined as any crypts with a clearly demarcated base lined with Paneth cells that also had at least 10 additional cell nuclei on either side of the crypt column beyond position 4, which is the generally accepted position of intestinal CBCs. If an intact crypt was identified, the number of positively staining cells in that individual crypt was recorded. Data was compiled using Microsoft Excel.

### Immunofluorescence (IF)

The small intestines were harvested, flushed with ice-cold PBS, cut into smaller fragments and fixed overnight in 10% buffered formalin. For cryopreservation, fixed intestines were next transferred to 30% sucrose in PBS overnight at 4 °C and stored in OCT at −80 °C. OCT sections (16-μm thick) were then permeabilized in 0.5% Triton (Sigma) and 0.2% Tween (Sigma #P1379) in PBS for 30 min, blocked in 2.5% BSA for 1 h and mounted in Mowiol mounting Media with DAPI (Invitrogen). Images were taken at 10X for crypt quantification, and 2.5X for representative full images using a Leica fluorescent microscope. For p53, Epcam and Clu-GFP staining, slides were incubated with antibodies to p53 protein (CM5) (Leica Biosystems), EpCAM (CD326) (G8.8) (118201; BioLegend) and GFP (Abcam #ab13970; 1:1000). To quantify p53 activation in the intestinal epithelia, all images were acquired at 63X on a Nikon Ti2 confocal microscope system. Images were processed in batch format in CellProfiler ^31^. DAPI counterstaining was used to create nuclear masks following measurement of p53 signal intensity in each nucleus, which was used to calculate frequency of cells showing p53 activation per image across all treatments.

For IF in intestinal organoids, organoids were grown on glass coverslips for 3-5 days after passaging. At the desired timepoint, organoids were fixed in 4% PFA (Electron Microscopy Sciences #15710) for 30 minutes, followed by permeabilization and blocking for 1 hour at room temperature in PBS with 0.5% Triton-X-100 (Sigma #X100-500ML)/0.2% Tween-20 (Sigma #P1379)/2% BSA (Sigma #A7906). Coverslips were incubated overnight at 4°C in primary antibodies targeted against p53 protein (CM5) (Leica Biosystems; 1:300) and GFP (Abcam #ab13970; 1:1000). Secondary DyLight antibodies (Fisher Scientific; 1:1000) were used. Sections were counterstained with DAPI (Sigma-Aldrich) for 30 minutes before mounting. Images were acquired using a Nikon Ti2 inverted confocal microscope platform.

### RNA in situ hybridization

RNA in situ hybridization (ISH) was performed with RNAscope technology (Advanced Cell Diagnostics). The probes to detect *Ly6a*, *Clu*, and *mKi67* mRNA were MmLy6a-427571, MmCLuC3-427891-C3 and Mmki67C2-416771, respectively. For chromogenic assay, ISH was conducted using RNAscope 2.5 HD Assay-BROWN as per the manufacturer’s protocol. For fluorescent assay, RNAscope Multiplex Fluorescent V2 Assay was used. Either the Opal Fluorophore reagent packs (Opal 520, Opal 570, Opal 690) (Akoya Biosciences) or the tyramide signal amplification (TSA) fluorophores (TSA Fluorescein, TSA Cyanine 3, TSA Cyanine 5) (Akoya Biosciences) were used for detection. Images were acquired using a Nikon Ti2 inverted confocal microscope platform.

### RNA ISH Image quantification

mKi67, Ly6a and Clu positive cells from RNA ISH tissue sections were quantified using QuPath v0.3.2 software ^32^. All cells were first detected based on DAPI staining using the *Cell detection* command. For each staining, a classifier was generated using the *Train object classifier* command. The three classifiers were then combined and run across all images. For each time point between 3-6 mice were harvested and total of 15-20 images were quantified for each intestinal swiss roll.

### Intestinal Crypt isolation and Single-cell suspension protocol

Small intestines were removed and around 15 cm of the jejunum was isolated and washed with ice-cold 1X HBSS (Cytiva, #SH3058801), opened longitudinally, and divided into three pieces of roughly equal length. To obtain the epithelial cells, intestinal fragments were incubated in epithelial cell solution (10 mM EDTA [Invitrogen, #15575020]-HBSS, 5 mM HEPES [Invitrogen, #15630080], 100 U/ml Pen-Strep [Invitrogen, #15140122], and 2% FBS [Life Technologies, #26140079]) for 30 min on ice with continuous gentle shaking. Tissue fragments were then transferred into fresh ice-cold HBSS. Vigorous shaking was used to isolate the epithelial fraction. Intestinal crypts were recovered by filtering the tissue suspension through a 70-μm mesh-cell-strainer. Three consecutive epithelial fractions were collected, pooled, and centrifugated for 10 minutes at 500 g. Epithelial cell pellets were resuspended in pre-warmed Trypsin-EDTA (Corning), and incubated 5 min at 37°C. Single cells were obtained by continuous pipetting for 15-20 minutes at room temperature (RT). Trypsin was quenched by the addition of 100% FBS and cells were washed twice with HBSS and finally filtered through a 40- μm mesh-cell-strainer.

### Flow Cytometry Cell Sorting and 10X Single-Cell RNA Sequencing Processing

Intestinal single cells were resuspended in fluorescence activated cell sorting (FACS) buffer (Advanced-DMEM/F-12 with 5% FBS) containing 0.1µg/mL of DAPI. For each sample, 1×10^5^ alive single cells were sorted in FACS buffer. After sorting, cells were washed twice with ice-cold PBS and counted again using an automated cell counter. 1×10^4^ cells for each sample were subjected to 10x Genomics single-cell isolation using the Chromium Next GEM Single Cell 3’ v3.1 Kit (10x Genomics) following the manufacturer’s recommendations. Single Cell 3’ Gene Expression libraries were sequenced using an Illumina NovaSeq 6000 platform.

### Single-cell RNA sequencing

#### Single-Cell RNA Seq Data Processing and Integration

The scRNA-seq data was processed using the Cell Ranger pipeline (10x Genomics) to demultiplex the FASTQ reads, align them to the mm10 Mouse genome, and generate gene-barcode expression matrices. Quality control of these expression matrices were conducted using the Seurat R package to retain cells with greater than 500 unique UMIs, more than 250 detected genes, a logarithmic ratio of detected genes per unique UMI above 0.8, and a proportion of counts in mitochondrial genes less than 10%. Additionally, only genes with non-zero counts in more than ten cells were kept, and doublet removal was performed using scDblFinder. Epithelial cells were first identified, and subset based on the expression of the epithelial gene marker Epcam. Identification of specific gene markers for all clusters was calculated using Seurat’s FindAllMarkers default settings. Five top enriched genes for each cluster were plotted using the DoHeatmap function from Seurat package. Epithelial cell clusters were then annotated based on the expression of canonical stem cell and differentiation markers previously described ^5, 33–35^. Enterocytes were further classified based on the expression of previously described villus zonation gene signatures ^9^. The *FeaturePlot* and *VlnPlot* functions were used to plot the expression of single genes in the UMAP and violin plots respectively. Differential gene expression was conducted using the default voom and limma-trend pipeline. Subsequent gene set enrichment analyses based on all differentially expressed genes were conducted with the Hallmark databases using the ClusterProfiler R package. To test how single-cell clusters identified in wild-type intestinal epithelia were altered upon p53 mutation, single-cell RNA-seq dataset generated from p53 mutant intestinal epithelia was first processed as above by using the same parameters that were applied to process the wild-type dataset. Next, FindTransferAnchors function implemented in Seurat was employed to compare the two datasets, where pre-calculated wild-type clustering was used as a reference map to infer single-cell clusters in the mutant dataset. MapQuery function was then used to generate a query map of mutant single cell clusters and plot the results.

### Statistical Analysis

GraphPad Prism (GraphPad Software) was used for the statistics in this study. To test statistical significance between samples from two different groups, two-tailed Student’s t-tests were used. When comparing several groups, a 2-way ANOVA test was used. For Kaplan-Meier curves, a Gehan-Breslow-Wilcoxon test was applied. All group data are represented by the mean and errors bars are the standard error of the mean (s.e.m).

