## Supplementary figures and images for "p53 promotes revival stem cells in the regenerating intestine after severe radiation injury"

### Figure S1-1

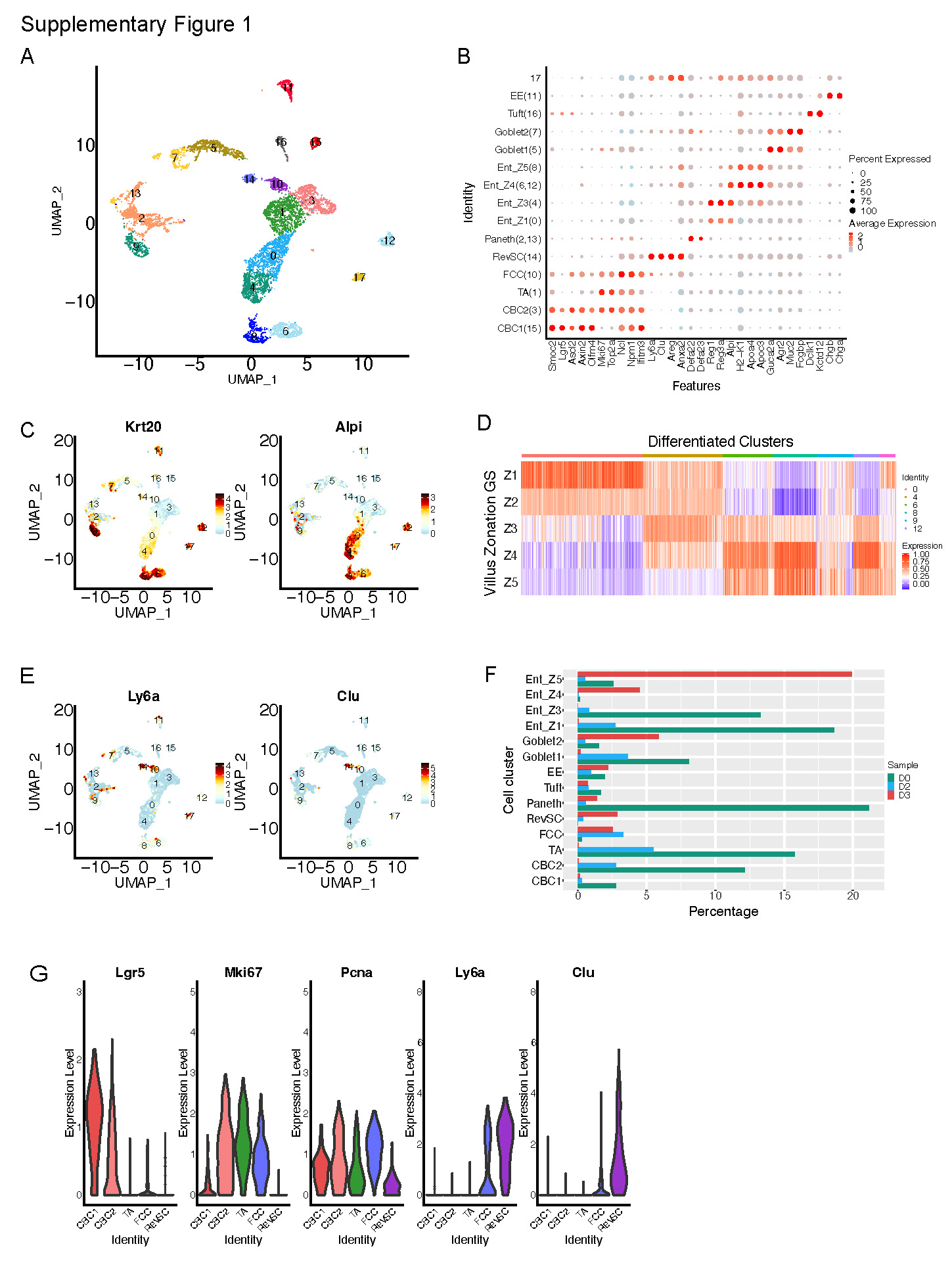

### Figure S1-2

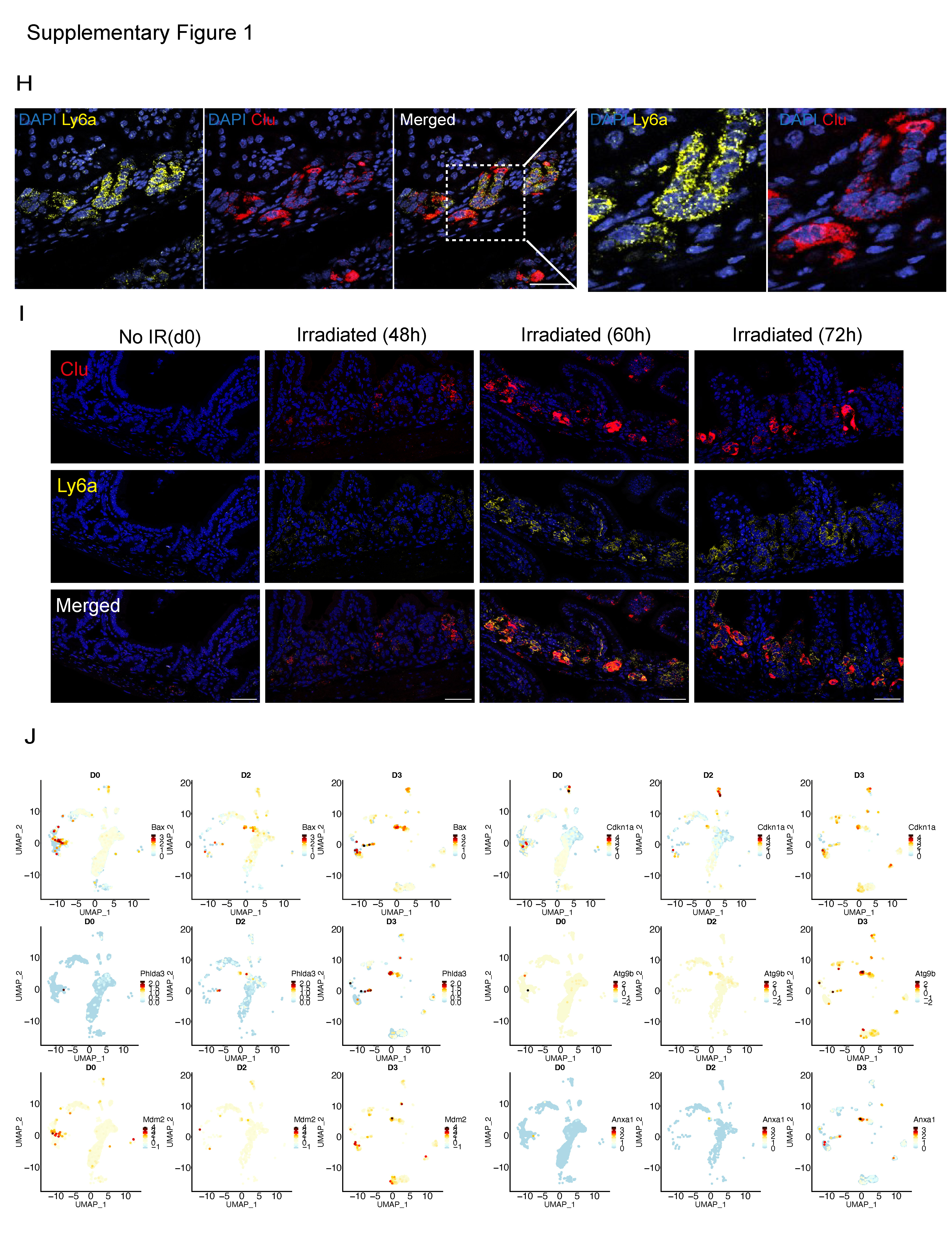

### Figure S2

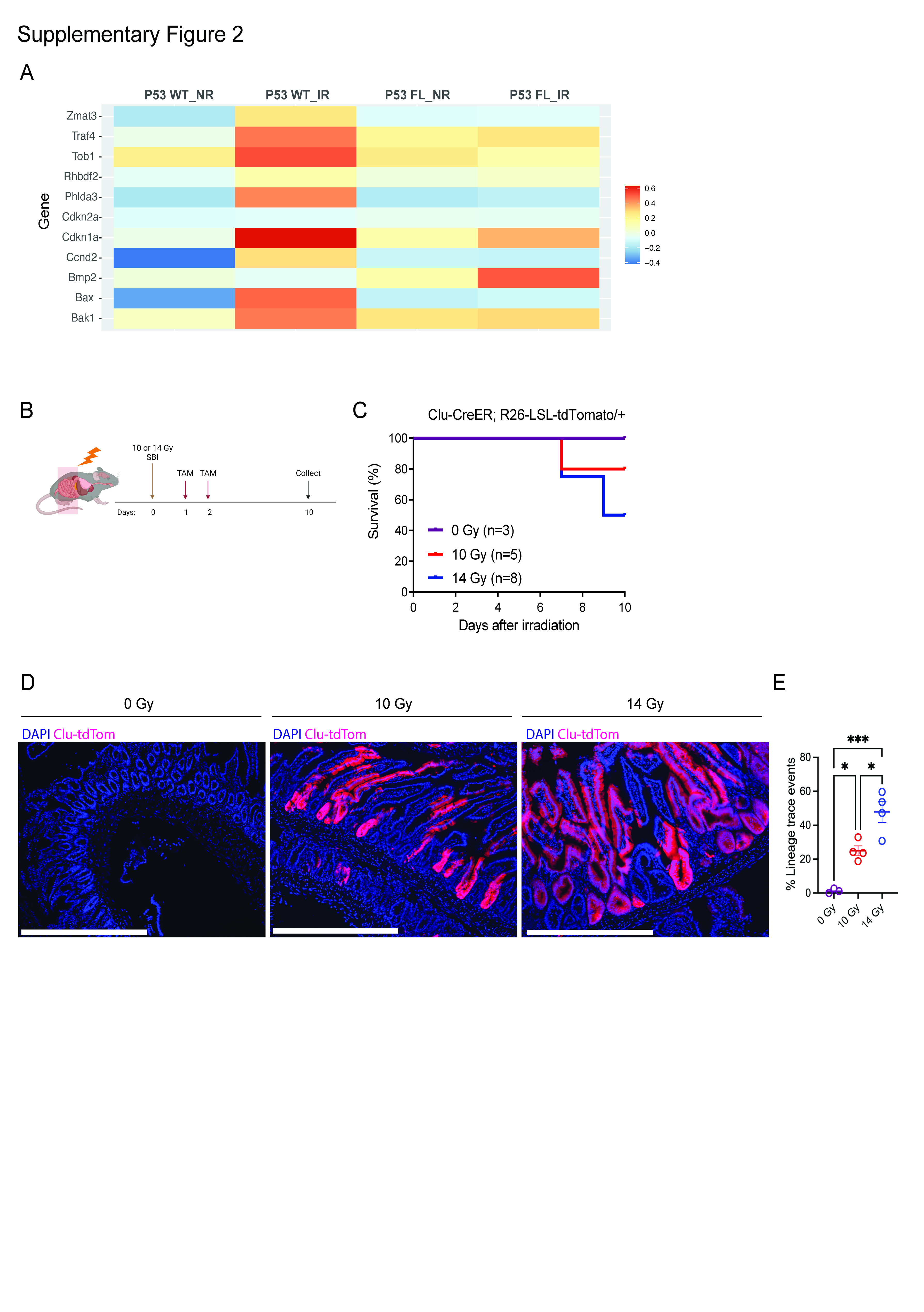

### Figure S3

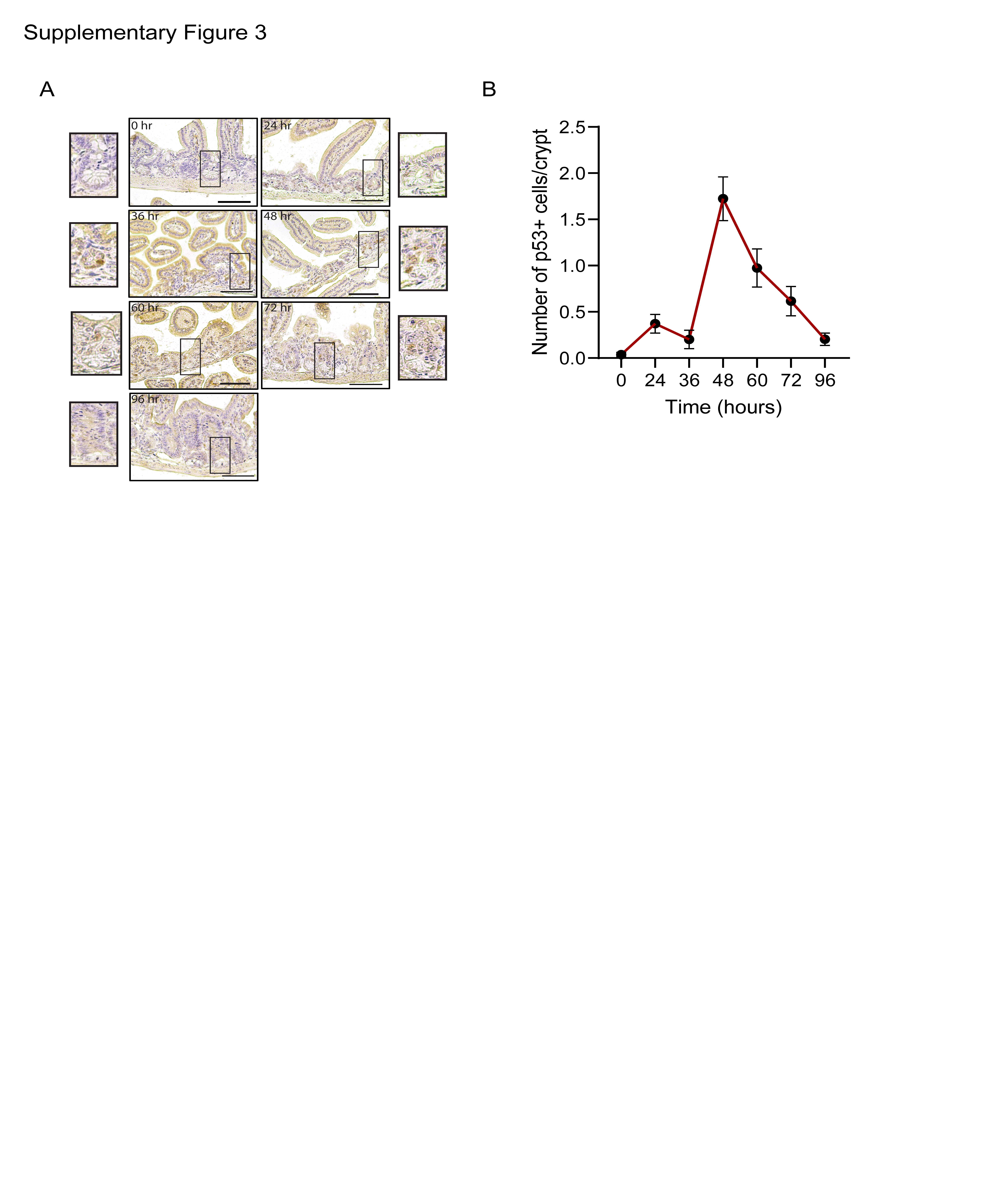

### Figure S4

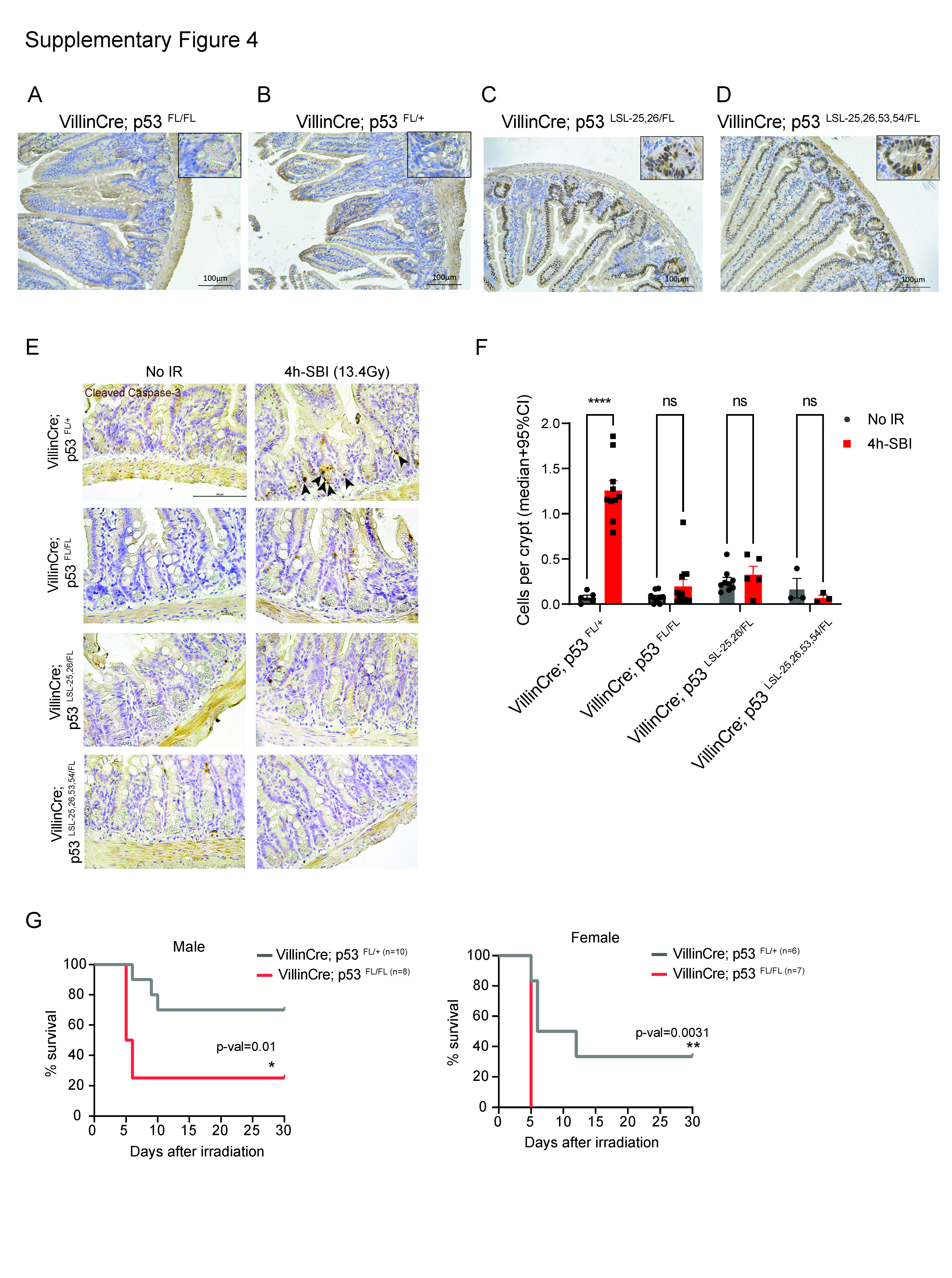
